# Deep proteomic analysis of Dnmt1 mutant/hypomorphic colorectal cancer cells reveals dys-regulation of Epithelial-Mesenchymal Transition and subcellular re-localization of Beta-Catenin

**DOI:** 10.1101/547737

**Authors:** Emily H Bowler, Alex Smith-Vidal, Alex Lester, Joseph Bell, Zhenghe Wang, Chris Bell, Yihua Wang, Nullin Divecha, Paul Skipp, Rob M. Ewing

## Abstract

**Background:** DNA methyltransferase I is the primary eukaryotic DNA methyltransferase engaged in maintenance of CpG DNA methylation patterns across the genome. Alteration of CpG methylation patterns and levels is a frequent and significant occurrence across many cancers, and targeted inhibition of Dnmt1 has become an approach of choice for select malignancies. There has been significant interest both in the methyltransferase activity as well as methylation-independent functions of Dnmt1. A previously generated hypomorphic allele of Dnmt1 in HCT116 colorectal cancer cells has become an important tool for understanding Dnmt1 function and how CpG methylation patterns are modulated across the genome. Colorectal cancer cells with the Dnmt1 hypomorphic allele carry a homozygous deletion of exons 3 to 5 of Dnmt1, resulting in greatly reduced Dnmt1 protein expression whilst still exhibiting a limited functional activity and methyltransferase ability. Although this cell model of reduced Dnmt1 levels and function have been used to study the downstream effects on the epigenome and transcriptome, the broader effects of the Dnmt1 hypomorph on the proteome and wider cell signalling are largely unknown.

**Results:** In this study, we used quantitative proteomic analysis of nuclear-enriched samples of HCT116 Dnmt1 hypomorph cells to identify signalling pathways and processes dysregulated in the hypomorph cells as compared to wild-type HCT116 cells. Unexpectedly, we observed a clear signature of increased expression of Epithelial-to-Mesenchymal (EMT) in Dnmt1 hypomorph cells. We also observed reduced expression and sub-cellular re-localization of Beta-Catenin in Dnmt1 hypomorph cells. Expression of wild-type Dnmt1 in hypomorph cells or knock-down of wild-type Dnmt1 did not recapitulate or rescue the observed protein profiles in Dnmt1 hypomorph cells suggesting that hypomorphic Dnmt1 causes changes not solely attributable

**Conclusions:** In summary we present the first comprehensive proteomic analysis of the widely studied Dnmt1 hypomorph colorectal cancer cells and identify redistribution of Dnmt1 and its interaction partner Beta-Catenin as well as the dysregulation of EMT related processes and signalling pathways related to the development of a cancer stem cell phenotype.

## INTRODUCTION

DNA methyltransferase 1 (Dnmt1) is the most abundant DNA methyltransferase in somatic cells [1-3], and mediates the fidelity of CpG methylation patterns in newly generated daughter cells [3, 4]. Loss of Dnmt1 is embryonically lethal with mouse embryonic stem cells lacking the catalytic function of Dnmt1 arresting prior to the 8-somite stage [2] and complete knockout of Dnmt1 in cultured cells results in G2/M cell cycle checkpoint arrest followed by apoptosis [2, 5]. Dnmt1 protein and its methyltransferase activity are required to maintain an undifferentiated and inexhaustible proliferative state in many stem cells and knockdown of Dnmt1 subsequently leads to premature differentiation of cells and eventually apoptosis as a result of loss of methylation markers [6]. Increased abundance of Dnmt1 protein has been shown in several cancer types, and typically correlates with tumour progression and altered genomic CpG methylation levels [7-10]. Dys-regulated DNA methylation activity results in cancer-specific methylation-dependent suppression of targeted gene expression, acting in a similar way to mutations that adversely affect tumour suppressor genes in developing tumour cells [4, 11, 12].

Increasing evidence indicates that Dnmt1 also has a range of DNA methyltransferase-independent functions. It has previously been shown that genes with changes in expression in response to Dnmt1 knockdown did not always contain CpG islands or show changes in their methylation status [1]. Through its interaction with Ubiquitin like with PHD and ring finger domains 1 (Uhrf1), Lysine acetyltransferase 5 (Tip60), and Histone deacetylase 1 (HDAC1), Dnmt1 can modulate acetylation, as well as suppress gene transcription as part of a regulatory complex with Chromodomain helicase DNA binding protein 4 (CHD4), and HDAC1 [13, 14]. Dnmt1 can function as a scaffold protein in transcription repression complexes independent of its methyltransferase catalytic domain, though its interaction with lysine demethylase 1A (LSD1). We previously showed that Dnmt1 can interact with Beta-Catenin in the nucleus of colorectal cancer cells leading to mutual stabilization of both proteins and increased CpG methylation and oncogenic Wnt signalling [15].

Due to the critical functions of Dnmt1, viable models of gene knockout and protein knockdown are limited. Chemical-mediated degradation of Dnmt1 is usually induced by treatment with the cytosine chemical analog 5-Azacitidine or its deoxy analog decitabine, which integrate into DNA as a substitute for cytosine, inhibiting Dnmt1 ability to donate its methyl group and resulting in Dnmt1 degradation [16, 17]. Dnmt1 mutant alleles encoding proteins with altered methyltransferase activity or reduced abundance have been of great interest in the field, allowing cells to be propagated and the function of Dnmt1 to be studied. [18] generated a series of Dnmt1 mutant cell models derived from the colorectal cancer cell line HCT116, although designated Dnmt1-/- cells. [19] further characterized these cell lines as expressing a lower abundance (20%) of truncated hypomorph protein (Dnmt1^Δ3-5^) approximately 17kDa smaller than the WT protein. This homozygous HCT116-Dnmt1^Δ3-5^ cell line has the DMAP1 interaction domain and the PCNA interaction domain deleted at the N-terminus by the insertion of a hygromycin resistance gene [18]. This mutant protein results in a 96% decrease in methyltransferase activity, although it retains its catalytic domain and DNA binding ability and genomic methylation levels are only decreased by approximately 20% [18-20]. These mutant cell lines have been used as models to study the effect of Dnmt1 on global methylation levels and the reactivation of CpG island promoter controlled gene expression [21]. In addition [22] found that HCT116-Dnmt1^Δ3-5^ cells showed reduced Ca2+-dependent cell adhesion, indicating dysregulation of E-Cadherin cell-cell adhesion junctions, this effect was independent of Dnmt3b signalling. Further characterization indicated a reduced abundance of E-Cadherin protein, and a re-localization of Beta-Catenin away from the plasma membrane, resulting in activated canonical Wnt signalling. This phenotype was specific to HCT116-Dnmt1^Δ3-5^ cells suggesting a DNA-methyltransferase-independent role for the N-terminal domain of Dnmt1 in cell-cell adhesion maintenance [22].

To investigate how the hypomorphic Dnmt1 allele impacts on cellular phenotype and on the cellular proteome we undertook a comprehensive proteomic analysis of HCT116-Dnmt1^Δ3-5^ cells. Analysis of signalling networks and pathways that were dysregulated in the HCT116-Dnmt1^Δ3-5^ nuclear proteomic profile identified alteration of the proteasome, cell adhesion junctions, cytoskeletal architecture and EMT were identified. We focused on further investigation of the very clear and significantly differential signature of EMT activation observed in the hypomorphic Dnmt1 cells and of the significantly altered profile of Beta-catenin subcellular re-localization. To understand whether altered expression of EMT related proteins is a consequence of reduced Dnmt1 protein levels or is a direct result of the mutant Dnmt1^Δ3-5^ protein, shRNA knockdowns of WT Dnmt1 were performed. These experiments showed that reducing Dnmt1 protein levels did not result in the induction of EMT marker proteins as observed in Dnmt1 hypomorphic cells. In addition, expression of wild-type Dnmt1 protein in hypomorphic Dnmt1 cells was not able to alter the EMT signature seen in these cells, suggesting that the EMT marker signature in Dnmt1 hypomorphic cells is a result of the presence of the hypomorphic protein rather than reduced protein abundance. In summary, we have identified multiple altered pathways and processes in a key model of Dnmt1 function in cancer cells, furthering our understanding and appreciation of the methylation-dependent and independent roles of this key protein.

## RESULTS

The HCT116-Dnmt1^Δ3-5^ cell-line originally developed by Rhee et al., (2000) [18] and characterised by Egger et al., (2006) [19] were grown in culture and Dnmt1 status of the hypomorph cell line was confirmed by PCR using primers specific to exons 1 and 6 of genomic WT Dnmt1 (Table 1). Amplification of the endogenous Dnmt1 in WT HCT16 cells and HCT116-Dnmt1^Δ3-5^ cells confirmed a product of reduced size in the HCT116-Dnmt1^Δ3-5^ cells as described by Egger et al., (2006) (Figure 1A and 1B). Further characterization of the endogenous protein was carried out by Western blot analysis (Figure 1C and 1D). HCT116-Dnmt1^Δ3-5^ cells showed a signal with reduced protein size and abundance compared to WT Dnmt1 in WT HCT116 cells, as previously shown by Egger et al., (2006) and we confirmed that two different anti-Dnmt1 antibodies gave the same result (Figures 1C and 1D). We also investigated two known Dnmt1 interaction partners, Usp7 [23] and Beta-Catenin [15]. No effect was observed on overall Usp7 protein abundance; however, there was an evident decrease in Beta-catenin abundance in the HCT116-Dnmt1^Δ3-5^ cell line (Figure 1C).

**Table 1.**
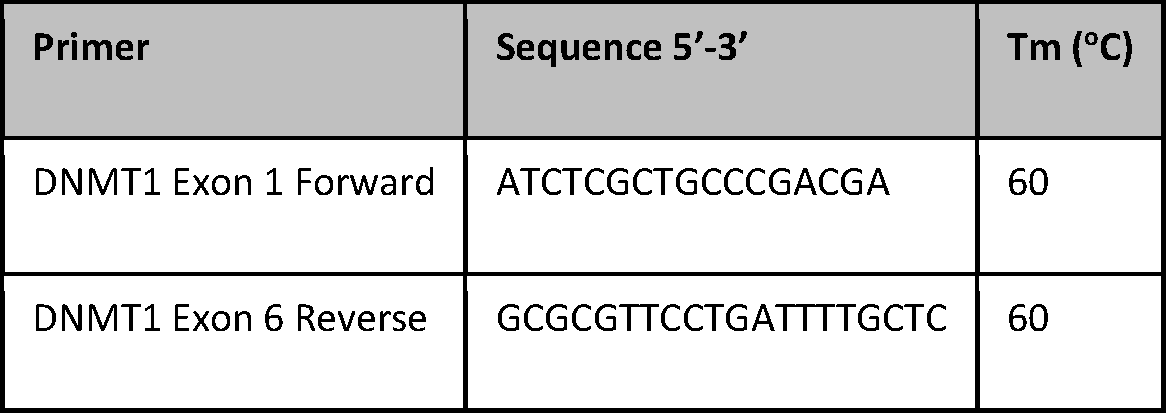
DNMT1 primers

**Figure 1.**
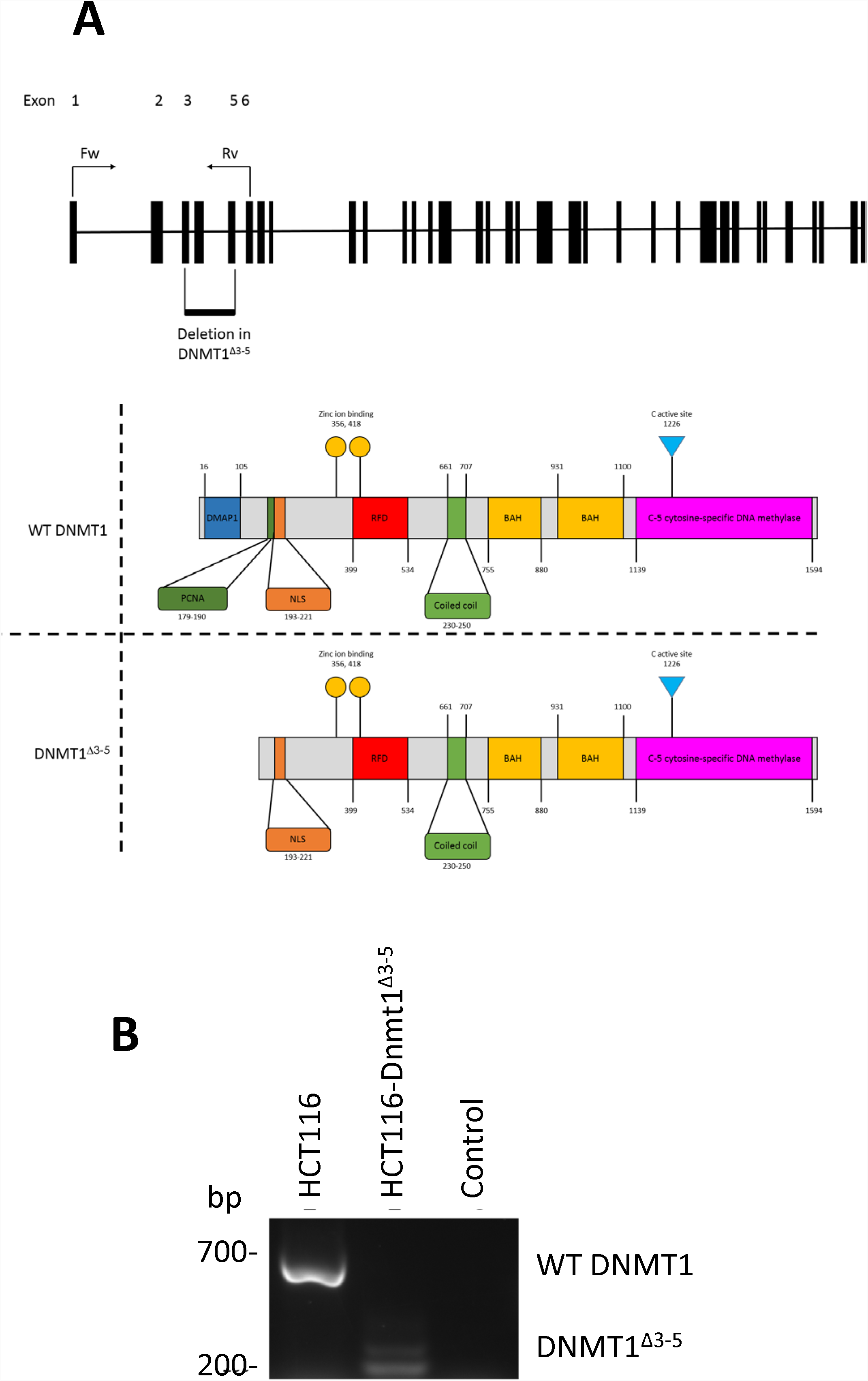
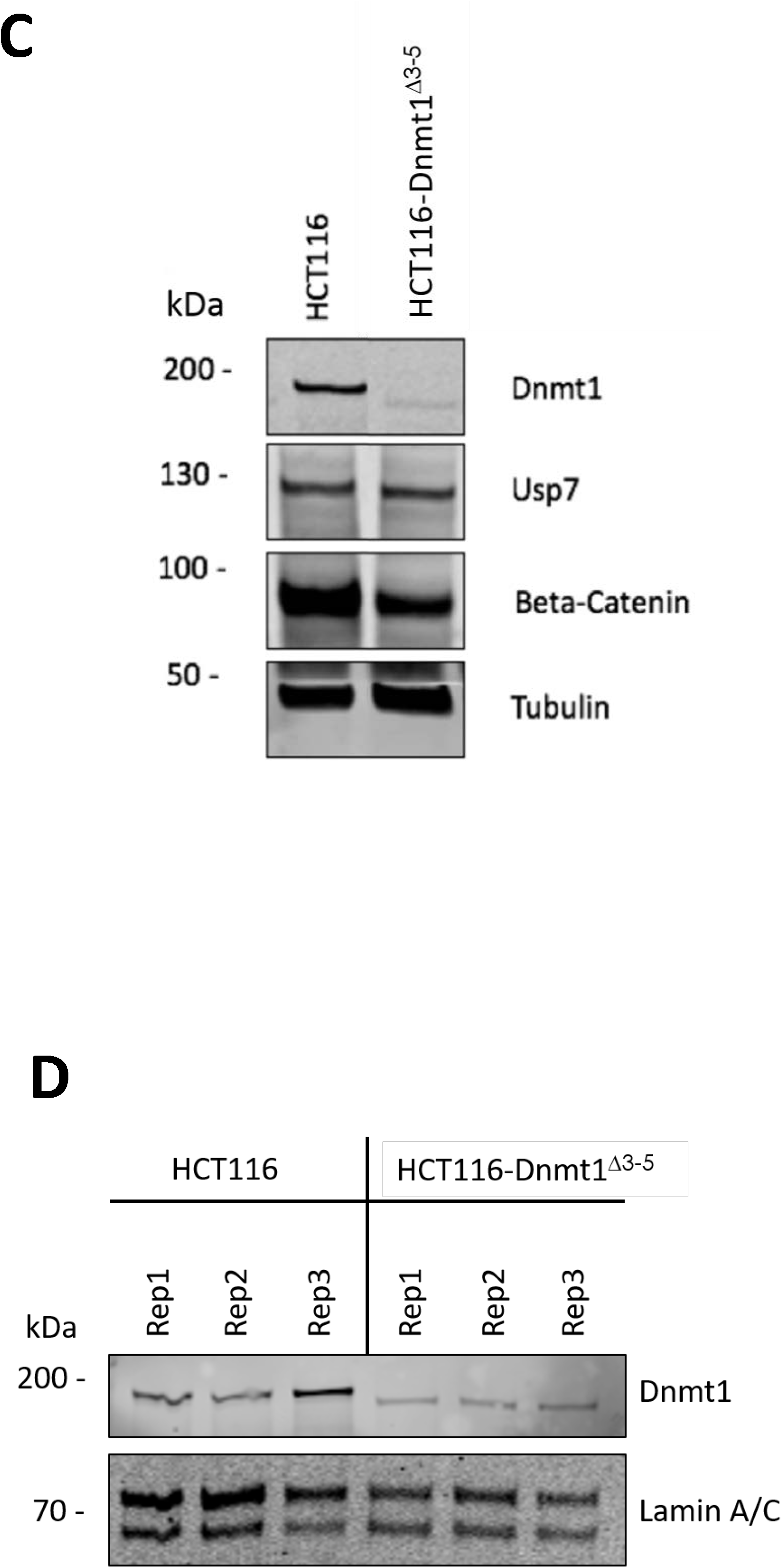
**A. Schematic diagram of DNMT1 exons and protein domains of HCT116 and HCT116-Dnmt1^Δ3-5^ cells.** Exon configuration of WT DNMT1, with annotation of primer sites used for analysis of DNMT1^Δ3-5^, and highlighted region of genomic deletion in this cell line. Protein domain schematic based on Pfam sites in Dnmt1 and Dnmt1^Δ3-5^. **B. Analysis of genomic Dnmt1 status in HCT116-Dnmt1^Δ3-5^ cells.** Polymerase chain reaction analysis of Dnmt1 in HCT116-Dnmt1^Δ3-5^ cells in comparison to the endogenous WT gene in HCT116 cells. mRNA was extracted from cells, converted to cDNA by reverse transcription and cDNA analysed by PCR using primers specific to DNMT1 exons 1 and 6 (table 2). PCR products were then analysed by gel electrophoresis resulting in a reduced DNMT1 product size in HCT116-Dnmt1^Δ3-5^ cells. **C. Western blot analysis of Dnmt1 and interaction partners in HCT116 and HCT116-Dnmt1^Δ3-5^ cells.** 15µg of whole protein lysates from HCT116 and HCT116-Dnmt1^Δ3-5^ cells were loaded onto a single phase 8% SDS gene and analysed by western blot analysis for protein expression of Dnmt1, Usp7, and Beta-Catenin. Tubulin protein abundance was used as a loading control. N=3. **D. Nuclear-enriched protein samples of HCT116 and HCT116-Dnmt1^Δ3-5^ cells prepared for mass spectrometry analysis of their nuclear proteomic profile.** 15µg of nuclear-enriched protein lysates from HCT116 and HCT116-Dnmt1^Δ3-5^ cells, three replicates of each, were loaded onto a single phase 8% SDS gene and analysed by western blot analysis for protein expression of Dnmt1. Lamin A/C protein abundance was used as a loading control for the nuclear-enriched lysate. These samples were then taken forward and prepared for nuclear proteomic analysis by data independent mass spectrometry analysis.

**Table 2.** Primary antibodies used in this study

| Antibody | Company | ID | Dilution | Host species |
| --- | --- | --- | --- | --- |
| Actin | Abcam | 8H10D10 | 1:1000 | Rabbit |
| B-Catenin | ThermoFisher | MA1-301 | 1:1000 | Mouse |
| δ-catenin | Cell signalling | 4989S | 1:1000 | Rabbit |
| Dnmt1 | Cell signalling | D63A6 | 1:1000 | Rabbit |
| E-Cadherin | BD Transduction Laboratories | 610182 | 1:1000 | Mouse |
| Histone H3 | Cell signalling | D18C8 | 1:1000 | Rabbit |
| Lamin A/C | Cell signalling | 2032S | 1:1000 | Rabbit |
| Lamin B1 | US Biological | L1220-21A | 1:1000 | Mouse |
| N-Cadherin | Cell signalling | D4R1H | 1:1000 | Rabbit |
| Slug | Cell signalling | C19G7 | 1:1000 | Rabbit |
| Snail | Cell signalling | L70G2 | 1:1000 | Mouse |
| Tubulin | Cell signalling | 2144S | 1:1000 | Rabbit |
| Twist | Cell signalling | 46702 | 1:1000 | Rabbit |
| Uhrf1 | Invitrogen | PA5-29884 | 1:1000 | Rabbit |
| Usp7 | Santa Cruz Biotechnology | Sc-30164 | 1:1000 | Rabbit |
| Vimentin | Cell signalling | D21H3 | 1:1000 | Rabbit |
| Zeb1 | Santa Cruz biotech | Sc-25388 | 1:1000 | Rabbit |

**Table 3.**
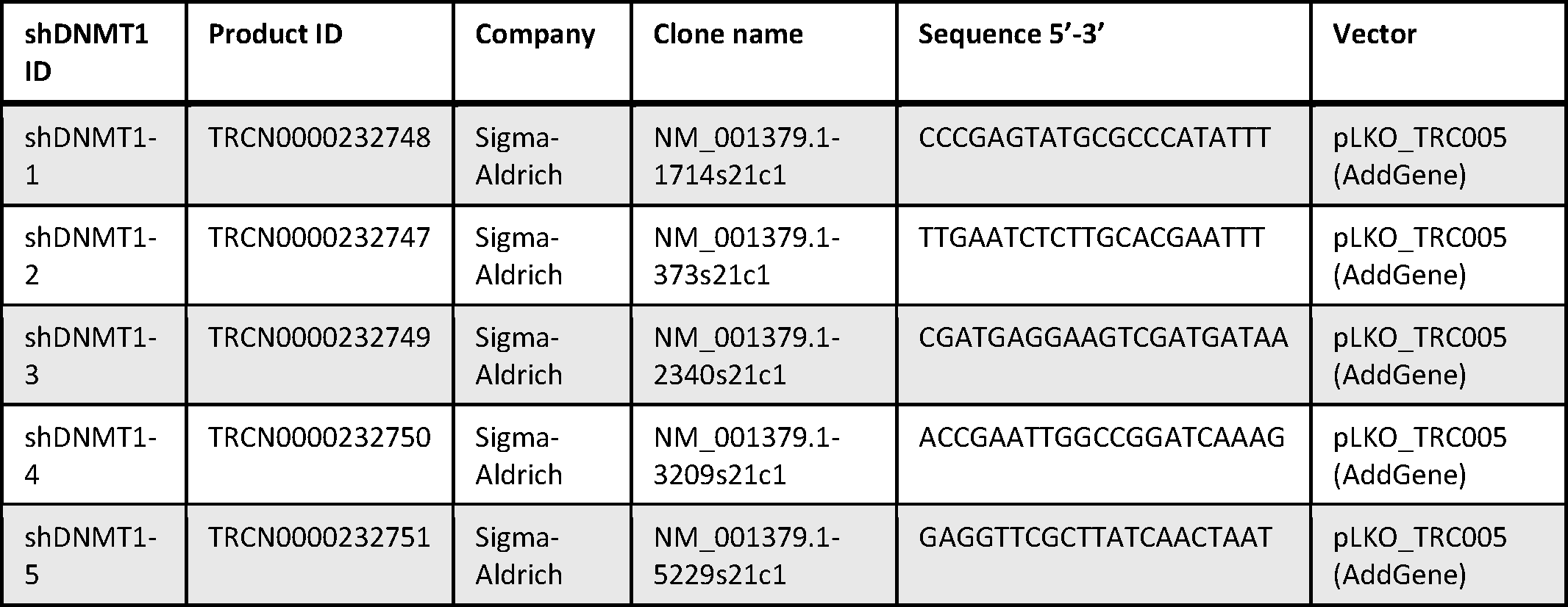
shRNA containing vectors used to knockdown DNMT1 expression Dnmt1 protein expression (derived from GenBank Accession: NM_001379)

To broadly characterize proteome-wide altered protein expression in HCT116-Dnmt1^Δ3-5^ cells we performed quantitative LC-MS/MS analysis on replicate nuclear samples from HCT116-Dnmt1^Δ3-5^ and HCT116 wild-type cells and characterised these same samples with Western blots as shown in Figure 1D. Data-independent quantitative mass spectrometry analysis identified a total of 3,678 proteins, of which 383 were found to be significantly differentially abundant (P value = <0.05). The sets of significantly differential proteins were analysed for enrichment of Gene Ontology (GO) terms. GO terms were ranked according to their significance in the HCT116 wild-type or HCT116-Dnmt1^Δ3-5^ cells and the most distinct terms are presented in Figures 2A and 2B. In particular, we noted the significant differences between the wild-type cells and hypomorph cells of their expression of proteasomal components (increased in wild-type cells), dys-regulation of metabolism (increased glyoxylate cycle, fatty acid and RNA metabolism in hypomorph cells) and cell-cell adhesion (increased expression of proteins in wild-type cells).

**Figure 2.**
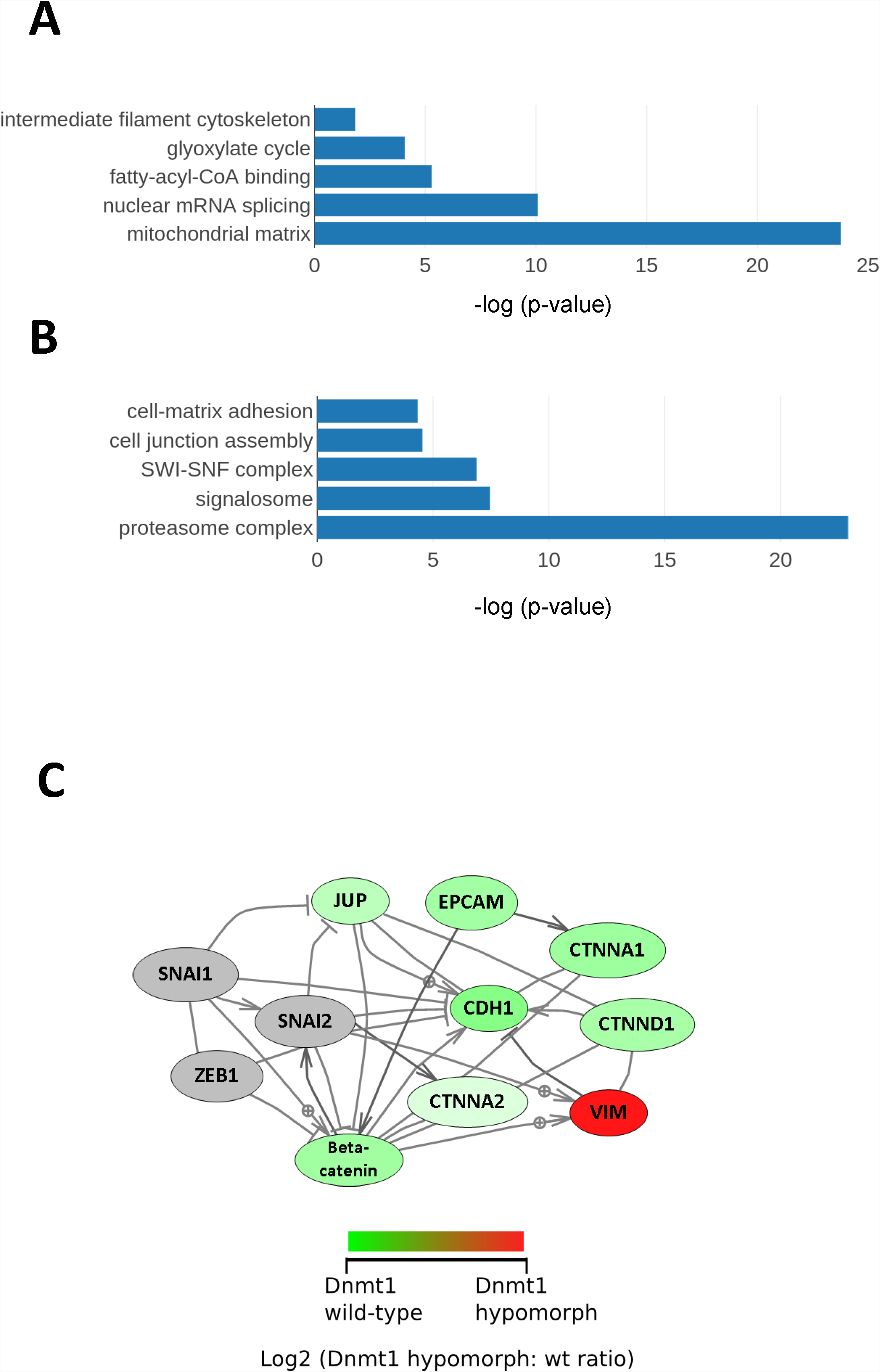
**A. Gene Ontology terms enriched in the proteome from HCT116-Dnmt1^Δ3-5^ cells.** Gene Ontology terms from the set of significantly (p<0.05) differential increased proteins in hypomorph cells. Selected GO terms where p-value <0.05 and the rank difference between hypomorph and wild-type GO terms are shown. **B. Gene Ontology terms enriched in the proteome from HCT116-Dnmt1^WT^ cells.** Gene Ontology terms from the set of significantly (p<0.05) differential increased proteins in wild-type cells. Selected GO terms where p-value <0.05 and the rank difference between wild-type and hypomorph GO terms are shown. **C. Network analysis of proteins with differential abundance in HCT116-Dnmt1^Δ3-5^ cells.** Selected differential protein interaction-networks are shown (Catenin-EMT network coloured according to mutant/wild-type abundance ratio. Nodes in grey indicate proteins not identified in the mass-spectrometry datasets but analysed by Western blot. Protein interaction networks were identified using Pathway Studio and String DB, using evidence-based directional interaction search settings. Networks were then extracted and manually pseudo-coloured to articulate differential abundance change in the nuclear proteomic profile of HCT116-Dnmt1^Δ3-5^ cells.

The protein with the largest increased abundance in HCT116-Dnmt1^Δ3-5^ cells was Vimentin (Log2 FC = 4.50), an intermediate filament protein with key roles in cytoskeletal architecture and cell signalling. In addition to this the top 50 proteins with increased differential abundance were enriched for proteins involved in amino acid metabolism, such as ACAT1, and DECR1, confirming the proposed dysregulation of amino acid metabolism in HCT116-Dnmt1^Δ3-5^ cells.

The protein with the largest decrease in abundance in HCT116-Dnmt1^Δ3-5^ nuclear proteome included KRT19, and KRT8 that are involved in cellular developmental processes, several members of the catenin protein family that enrich for the dysregulation of cell adhesion signalling, and members of the proteasome (prosoma, macropain) family that constitute proteosomal protein complexes. Large differential abundance changes in these proteins confirm the proposed dysregulation of protein ubiquitination-mediated degradation, cell adhesion, and cellular development as identified in GO biological process analysis.

As shown in Figure 2C, we identified a network of proteins that co-complex at the cell membrane forming cell-cell adhesion contacts via adherens junctions. Beta-catenin has previously shown to be an interaction partner of Dnmt1 in a mutual stabilizing interaction [15]. Alpha-Catenin, Delta-Catenin, and E-Cadherin form adherens junctions, connecting intracellular actin filaments to adjacent cells as one of the primary forms of cell-cell contacts in epithelial cells. All of the proteins in this complex exhibited decreased protein abundance in the nuclear proteomic profile of HCT116-Dnmt1^Δ3-5^ cells. The change in abundance of Delta-Catenin was confirmed using western blot analysis of whole cell and nuclear-enriched protein lysates validating that protein abundance levels are lower in HCT116-Dnmt1^Δ3-5^ cells.

Three myosin proteins were found to be dysregulated in HCT116-Dnmt1^Δ3-5^ cells, that regulate cytoskeletal architecture, migration and cell polarity [24]. In addition, these have been shown to interact with Arpc5l which facilitates actin polymerization, and POTEE which has been found up regulated in numerous cancer tissues and cell lines [25]. Taken together this interaction network regulates cellular architecture and structure, dysregulation of which is a key hallmark of tumourigenesis [26]

We observed dramatically altered expression of Vimentin (UniProt: P08670) between the HCT116 wild-type and HCT116-Dnmt1^Δ3-5^ cells and the fold-change of this protein was the largest statistically significant change observed across the complete dataset (log2 fold change=4.50; p-value = 1 x 10-3). Western blots of nuclear-enriched and whole cell lysates of key EMT markers are shown in Figure 3A and 3B, and corresponding ratios and p-values from the mass-spectrometry data are shown in Figure 3. Western blot analysis indicated that there was only low levels of Vimentin protein present in HCT116 cells, and significant expression of Vimentin in both the nuclear-enriched and whole cell protein samples extracted from HCT116-Dnmt1^Δ3-5^ cells, in agreement with a previous study [22]. Vimentin protein levels have previously reported to be low in WT HCT116, with western blot analysis with some antibodies producing negligible protein bands [27]. Vimentin is an intermediate filament protein, regulating cell motility, organelle and membrane-associated protein organization [25]. Significantly increased Vimentin expression is reminiscent of the typical phenotype shown by the process of epithelial to mesenchymal transition (EMT) and we next performed Western blot analysis of other key EMT protein markers. HCT116-Dnmt1^Δ3-5^ cells showed substantially decreased E-cadherin protein abundance in both nuclear and whole cell protein extracts when compared to HCT116 cells (Figure 3A) although N-cadherin protein abundance was unaltered (and not detected in the mass-spectrometry experiments). The Western blots confirmed the mass-spectrometry data, whereby no peptides corresponding to E-cadherin were detected in samples from the HCT116-Dnmt1^Δ3-5^ cells.

**Figure 3.**
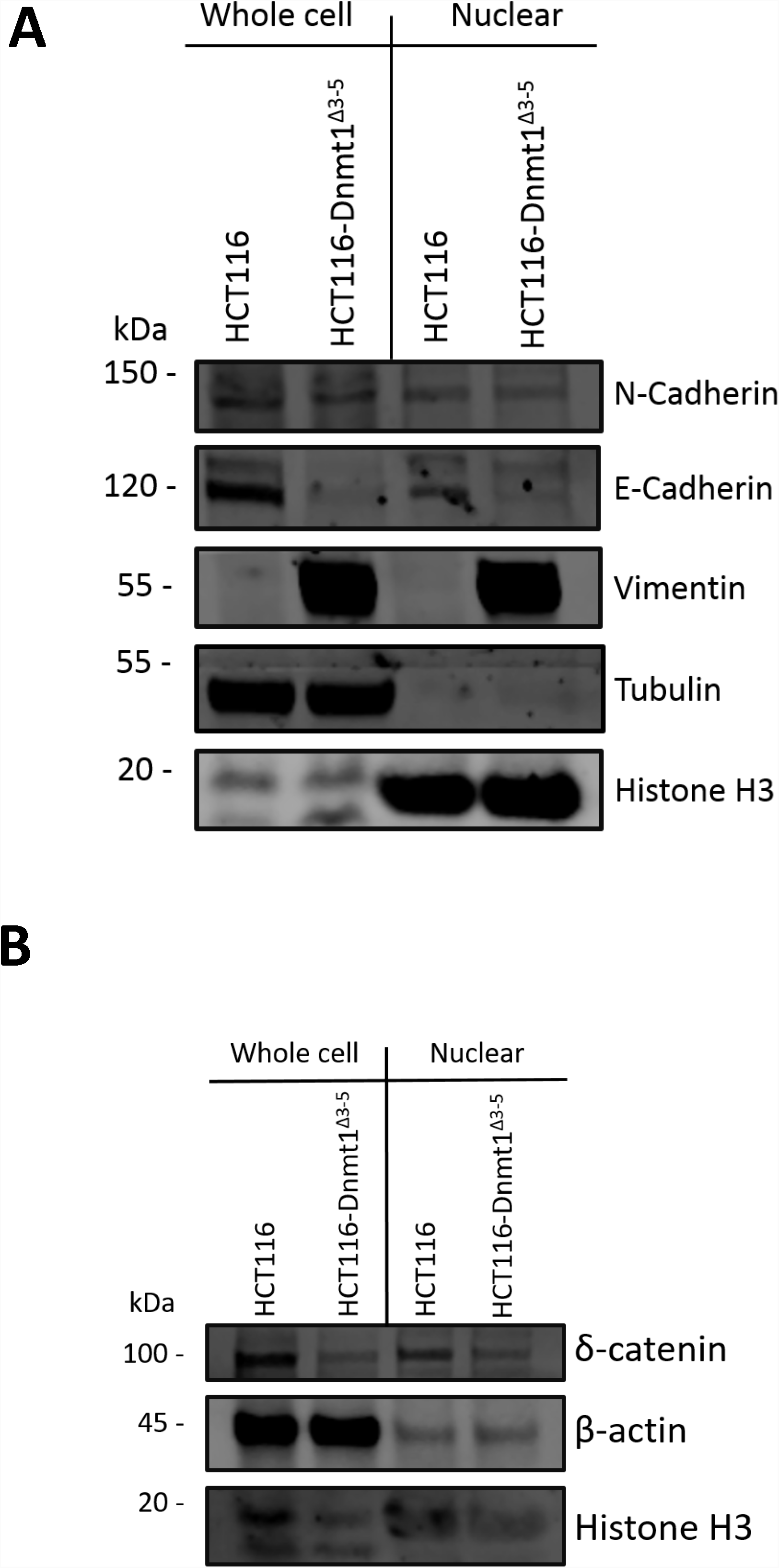
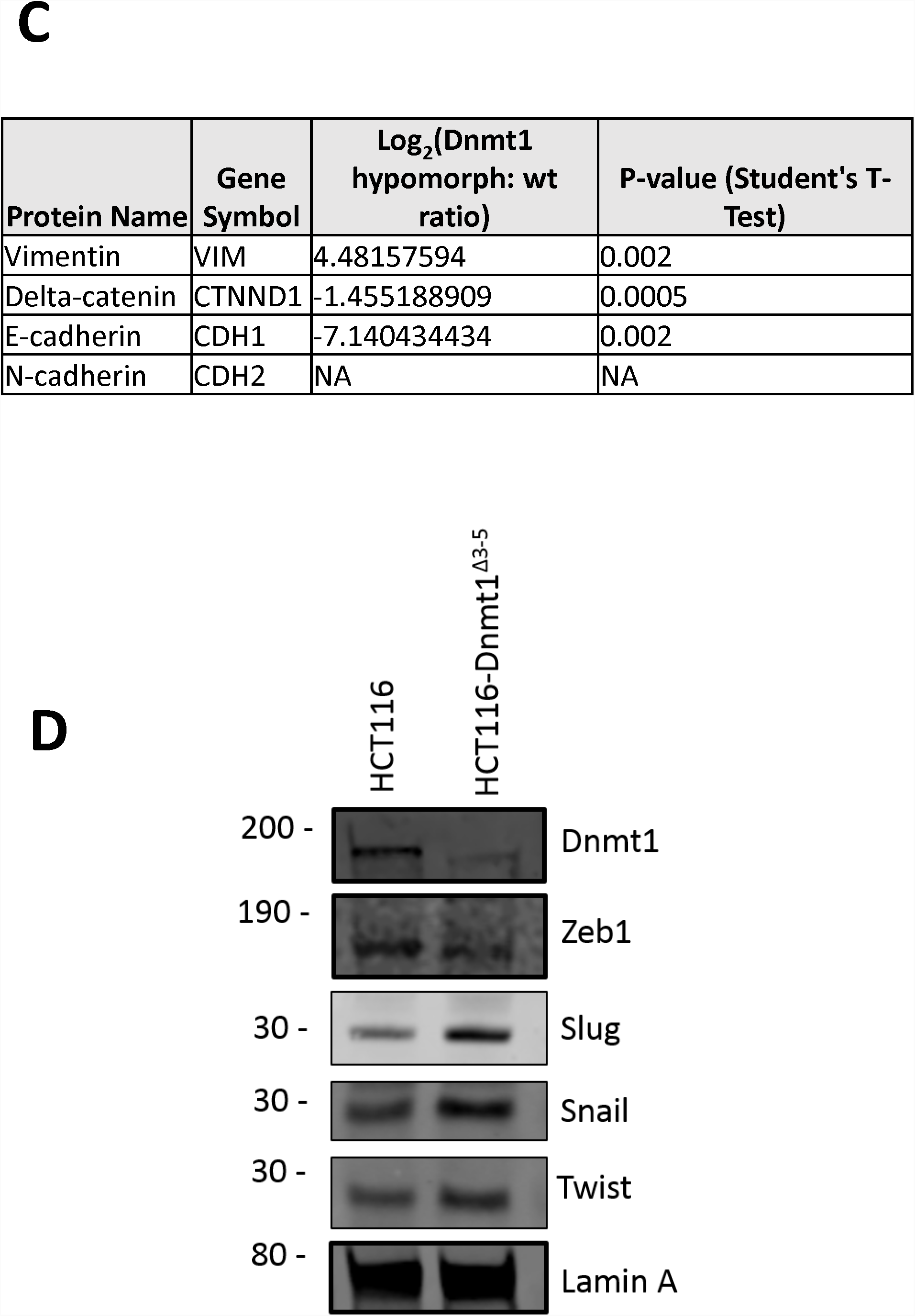
**A. Analysis of nuclear enriched and whole cell protein lysates for protein abundance of EMT markers.** 15µg of nuclear-enriched or whole cell protein lysates from HCT116 and HCT116-Dnmt1^Δ3-5^ cells were loaded onto a single phase 8% SDS gene and analysed by western blot analysis for protein expression of Vimentin, N-Cadherin, and E-Cadherin. Tubulin protein abundance was used as a loading control for the whole cell protein lysate and Histone H3 protein abundance was used as a loading control for the nuclear-enriched lysate. N=3. **B. Analysis of nuclear enriched and whole cell protein lysates for protein abundance of Delta-Catenin.** 15µg of nuclear-enriched or whole cell protein lysates from HCT116 and HCT116-Dnmt1^Δ3-5^ cells were loaded onto a single phase 8% SDS gene and analysed by western blot analysis for protein expression of Delta-Catenin. ß-Actin protein abundance was used as a loading control for the whole cell protein lysate and Histone H3 protein abundance was used as a loading control for the nuclear-enriched lysate. N=3. **C. Corresponding mass spectrometry protein quantification of Dnmt1 and key EMT protein markers. D. Western blot analysis of transcription-factors associated with EMT.** 15µg of nuclear-enriched protein lysates from HCT116 and HCT116-Dnmt1^Δ3-5^ cells were loaded onto a single phase 8% SDS gene and analysed by western blot analysis for protein abundance of Dnmt1, Zeb1, Slug, Snail, and Twist. Lamin A protein abundance was used as a loading control. N=3.

Following identification of dysregulation of key phenotypic EMT markers, we investigated the EMT-associated transcription factors Slug, Snail, Zeb1 and Twist (Figure 3D). Western blot analysis of these transcription factors showed that Slug, Snail and Twist are clearly elevated in HCT116-Dnmt1^Δ3-5^ cells compared to HCT116 wild-type cells, suggesting that activation of these transcription factors may be the drivers of altered E-cadherin and Vimentin expression. Slug and Snail have been shown to induce EMT signalling in TGFß treated CRC cell lines in tandem with Zeb1 and Twist signalling, however these transcription factors can act independently of one another to induce EMT transformation [27]. Increased protein expression and nuclear localization of Snail mediates the cadherin switch, resulting in the EMT phenotype of decreased E-Cadherin and increase N-Cadherin abundance [28]. E-Cadherin transcription can also be repressed by the binding of Slug, Zeb1, and Twist to its promoter, resulting in decreased protein abundance, these transcription factors also promote the increase in protein abundance of N-Cadherin, facilitating the progression of EMT [29]. We have therefore shown that an increase in Slug and Snail protein abundance in HCT116-Dnmt1^Δ3-5^ cells may drive the partial cadherin switch between E-Cadherin and N-Cadherin that is evident in these cell lines.

Beta-Catenin is another important driver of EMT and Vimentin expression, for which we observed significantly differential abundance in the mass-spectrometry data (log2 fold decrease=-2.2 in HCT116-Dnmt1^Δ3-5^; p-value=1 ×10-3) and by Western blot (Figure 1C). Beta-catenin is also a nuclear interaction partner of Dnmt1 in wild-type HCT116 cells [15] and we therefore analysed the subcellular distributions of Dnmt1 and Beta-catenin as shown in Figure 4A. Nuclear and cytosolic protein lysates from HCT116-Dnmt1^Δ3-5^ and wild-type cells were analysed and as expected Dnmt1 protein found to be significantly enriched in the nuclear samples [30]. Interestingly a similar ratio of Dnmt1 protein localization was not observed in HCT116-Dnmt1^Δ3-5^ cells, although there is an overall lower protein expression of Dnmt1^Δ3-5^ in these cells, there was approximately the same amount of protein present in both the nuclear enriched and cytosol enriched protein samples. This would suggest that even though the Dnmt1^Δ3-5^ protein retains its nuclear localization signal the distribution of the protein throughout the cellular compartments is modified in response to the loss of exons 3 to 5. To further investigate this finding, we used immunolocalization and confocal microscopy to visualize Dnmt1 and Beta-catenin sub-cellular distribution as shown in Figure 4B. A lower abundance of Dnmt1 protein was observed in HCT116-Dnmt1^Δ3-5^ cells, as well as a more diffuse localization of Dnmt1 in this cell line. Co-localization analysis of the confocal data indicated that Dnmt1 and Beta-Catenin co-localize within the nucleus of both cell lines, although there is a larger amount of ‘free’ Dnmt1 protein present in the nucleus of the HCT116 cell line. In addition to this, co-localization analysis of the HCT116-Dnmt1^Δ3-5^ cell line indicates that Dnmt1 present in the cytosol of these cell lines also co-localizes with Beta-Catenin, indicating that this interaction is not exclusive to the nucleus in this cell line. In alignment with the Western blot analyses, immunofluorescent imaging indicated that Beta-Catenin distribution was more diffuse throughout the HCT116-Dnmt1^Δ3-5^ cells, with a larger proportion of the protein present in the cytoplasm compared to the WT HCT116 cell line.

**Figure 4.**
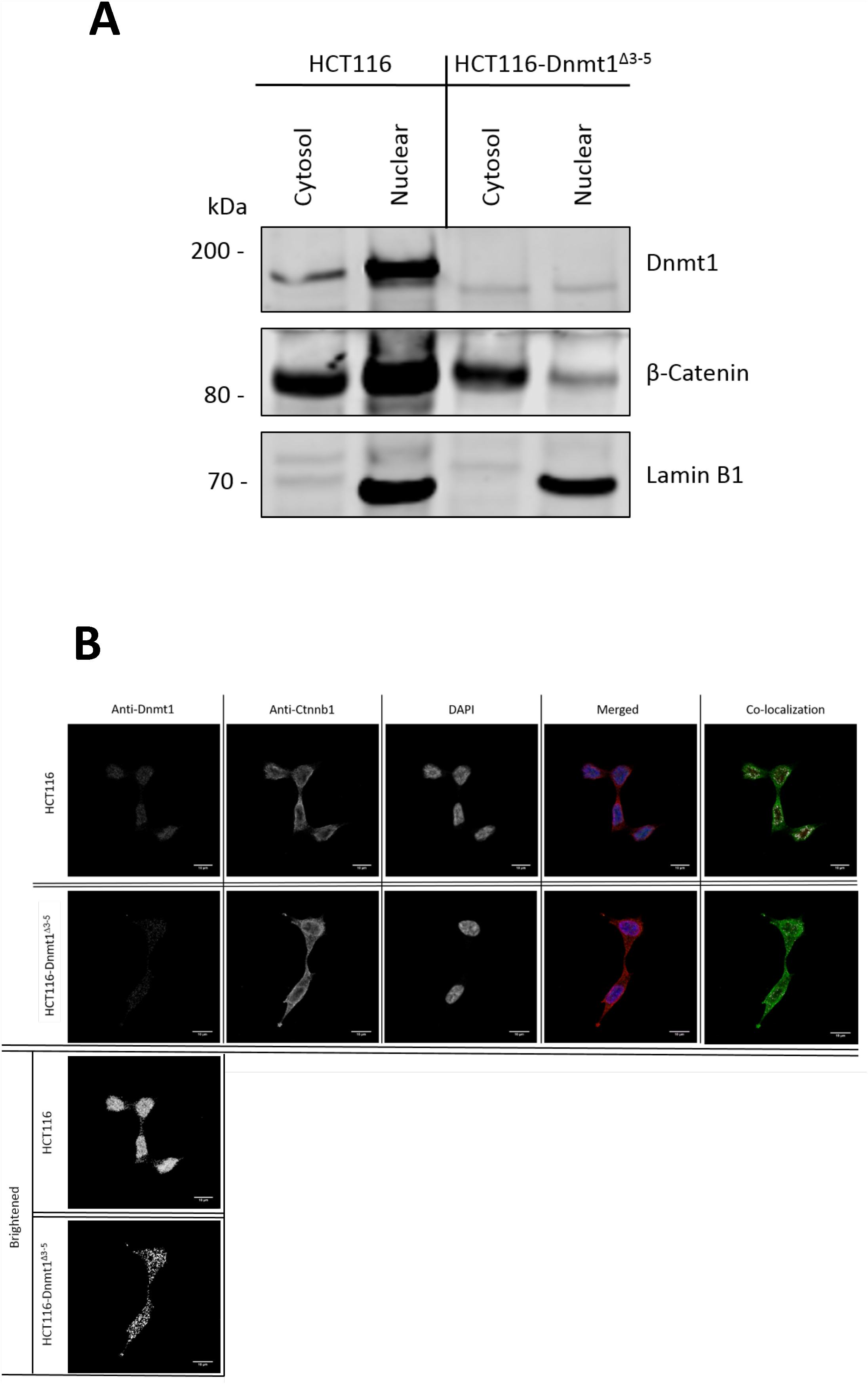
**A. Analysis of nuclear and cytosolic enriched protein lysates for Dnmt1 and Beta-Catenin localization.** 15µg of nuclear-enriched or cytosol-enriched protein lysates from HCT116 and HCT116-Dnmt1^Δ3-5^ cells were loaded onto a single phase 8% SDS gene and analysed by western blot analysis for protein expression of Dnmt1 and Beta-Catenin. Lamin B1 protein abundance was used as a loading control for the nuclear-enriched lysate. N=3. **B. Immunofluorescence analysis of Beta-Catenin and Dnmt1 localization in HCT116 and HCT116-Dnmt1^Δ3-5^ cells.** Localization of Dnmt1 and Beta-Catenin protein was assessed using fluorophore conjugated secondary antibodies and DAPI fixing agent. Channels are presented separately in addition to merged. Co-localization of Dnmt1 and Beta-Catenin was analysed using FIJI and MBF plugin, result of which is displayed in the ‘co-localization’ panel where areas of co-localization of Dnmt1 (red) and Beta-catenin (green) are highlighted in white.

We next investigated whether the reduced levels of Dnmt1 protein in HCT116-Dnmt1^Δ3-5^ cells was associated with the observed induction of EMT protein markers. Knock-down of Dnmt1 protein in wild-type HCT116 cells was achieved following transfection with 5 individual shRNA constructs as well as a mix of all 5 of these shRNA constructs, using a transfected scrambled shRNA construct as a control. Samples were analysed by western blot with WT HCT116 and HCT116-Dnmt1^Δ3-5^ protein samples as reference controls and a GFP plasmid transfected cell line as an independent transfection control. Western blots were used to determine protein abundance of the EMT markers Vimentin, E-Cadherin, and N-Cadherin. All shRNA constructs resulted in a decreased protein abundance of Dnmt1, apart from construct 1 (shDNMT1-1), the transfection protocol did not affect Dnmt1 protein abundance levels, as shown by the shDNMT1 (-ve) construct and GFP plasmid transfected HCT116 cell line (Figure 5A). Reduction of Dnmt1 by shRNA knockdown did not increase Vimentin expression in WT HCT116 cell lines. Similarly, no alterations to either E-Cadherin (Figure 5B) or N-Cadherin (Figure 5C) protein abundance by the shRNA-mediated knock-down of Dnmt1 protein expression were observed. We also tested whether expression of full-length Dnmt1 protein in HCT116-Dnmt1^Δ3-5^cells would result in decreased Vimentin expression seen in HCT116 wild-type cells (Figure 5D). Expression of the full length DNMT1 construct was identifiable in the HCT116-Dnmt1^Δ3-5^ cells on the western blot as a separate band with increased size, comparable to that shown in WT HCT116 cells. Protein samples from the Dnmt1 recovery experiment did not show a reduction in Vimentin protein abundance when compared to HCT116-Dnmt1^Δ3-5^ cells. Similar rescue experiments by [22] indicate that E-Cadherin expression can be reversed following expression of exogenous WT Dnmt1, however this paper did not stipulate the duration of WT Dnmt1 expression required to show this, nor did they investigate the effect of WT Dnmt1 on the ability to repress Vimentin protein expression.

**Figure 5.**
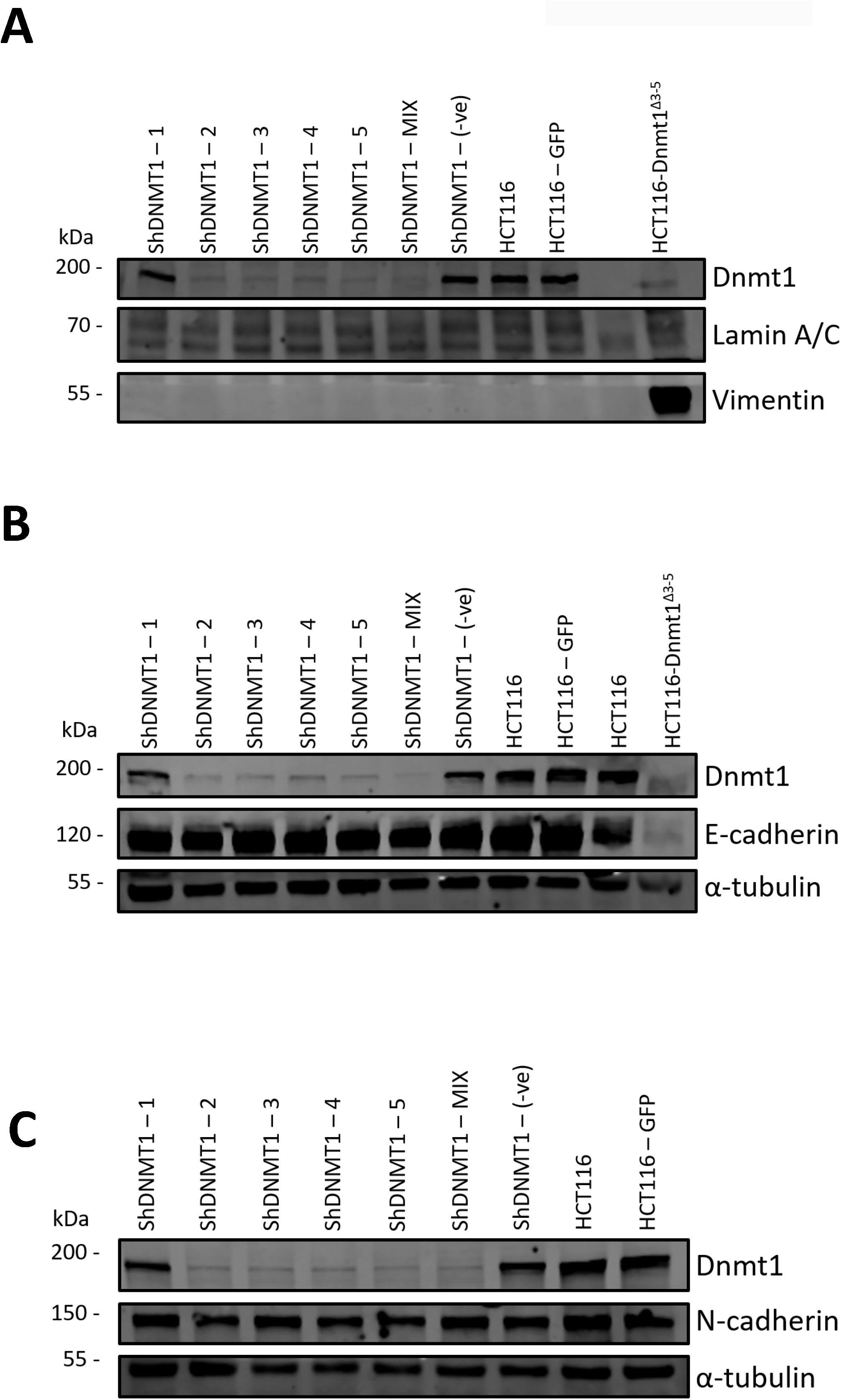
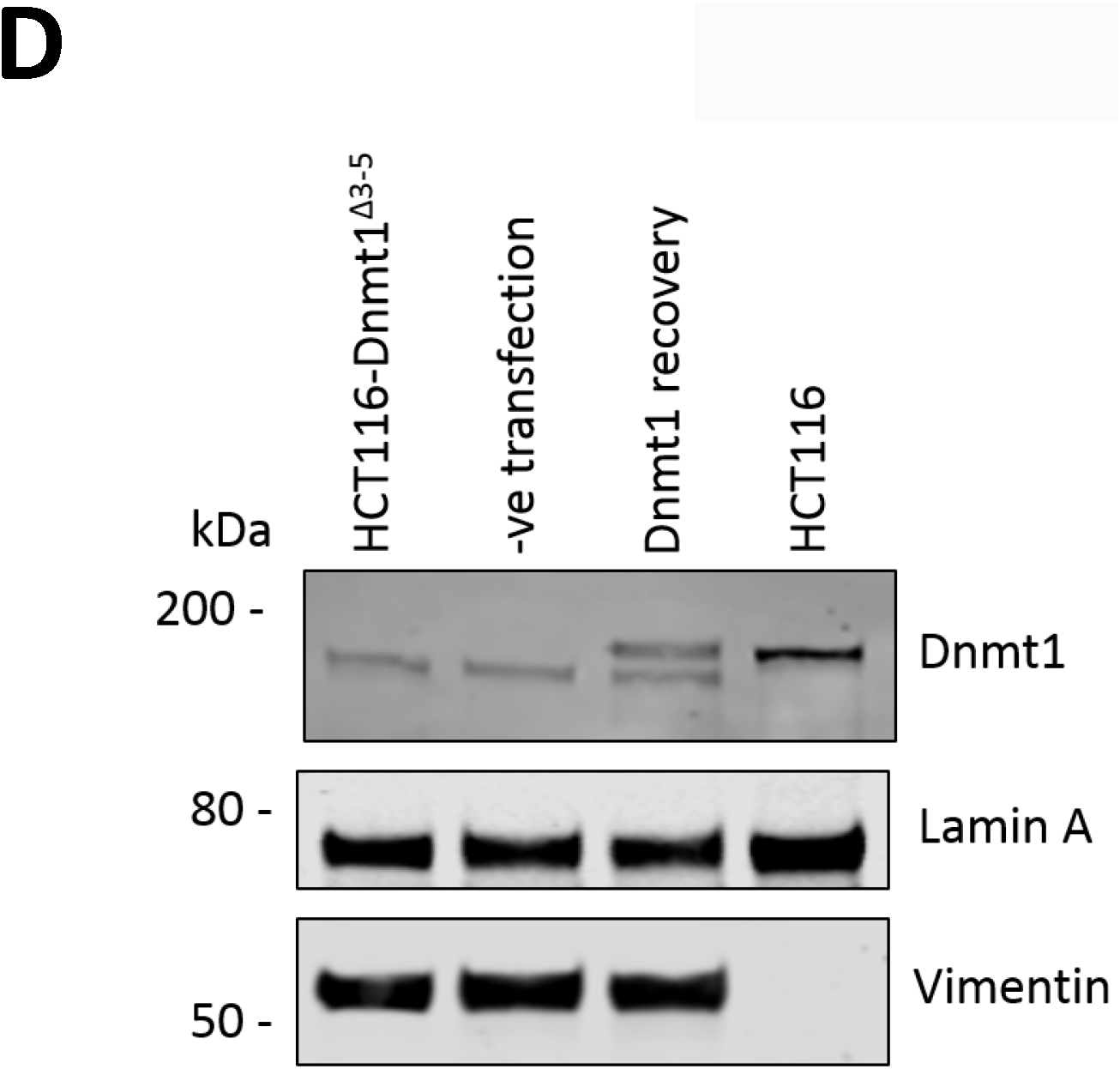
**A. Dnmt1 shRNA knockdown analysis on Vimentin protein abundance.** 15µg of whole cell protein lysates from shRNA knockdown HCT116 cells, HCT116-GFP controls cells and HCT116-Dnmt1^Δ3-5^ cells were loaded onto a single phase 8% SDS gene and analysed by western blot analysis for protein expression of Vimentin. A-Tubulin and Lamin A/C protein abundance were used as a loading control. N=3. **B. Dnmt1 shRNA knockdown analysis on E-Cadherin protein abundance.** 15µg of whole cell protein lysates from shRNA knockdown HCT116 cells, HCT116-GFP controls cells and HCT116-Dnmt1^Δ3-5^ cells were loaded onto a single phase 8% SDS gene and analysed by western blot analysis for protein expression of E-Cadherin. A-Tubulin protein abundance was used as a loading control. N=3. **C. Dnmt1 shRNA knockdown analysis on N-Cadherin protein abundance.** 15µg of whole cell protein lysates from shRNA knockdown HCT116 cells, HCT116-GFP controls cells and HCT116-Dnmt1^Δ3-5^ cells were loaded onto a single phase 8% SDS gene and analysed by western blot analysis for protein expression of N-Cadherin. A-Tubulin protein abundance was used as a loading control. N=3. **D. The effect of wild type Dnmt1 protein expression rescue in HCT116-Dnmt1^Δ3-5^ cells on Vimentin protein abundance.** 15µg of whole cell protein lysates from HCT116 cells, HCT116-Dnmt1^Δ3-5^ cells, and HCT116-Dnmt1^Δ3-5^ cells transfected with either a full length WT Dnmt1 expression vector or an empty control were loaded onto a single phase 8% SDS gene and analysed by western blot analysis for protein expression of Dnmt1. Tubulin protein abundance was used as a loading control. N=2.

## DISCUSSION

HCT116-Dnmt1^Δ3-5^ hypomorph cells are a widely used model of altered CpG methylation in colorectal cancer cells. These cells exhibit reduced catalytic methyltransferase activity and show altered patterns of CpG methylation [18, 19]. Studies of Dnmt1 by knockdown or knockout methods are typically limited due to the criticality of the protein, therefore this HCT116-derived cellular model with reduced mutant Dnmt1 protein abundance and activity has allowed the elucidation of downstream Dnmt1 targets typical of its role as a methyltransferase [21]. Studies of the effects of this mutant Dnmt1 on protein signalling networks however is less well understood. We present the first comprehensive proteomic analysis of Dnmt1 hypomorph cells and identify key protein and pathway alterations in these cells. Here we show multiple signalling pathways are altered in the nucleus of HCT116-Dnmt1^Δ3-5^ cells, which include changes in cell-cell adhesion, proteasome complex organization, and cellular architecture. Key regulators of the EMT process were amongst the most differentially expressed proteins in the dataset.

Further investigation of the EMT process shown to be dysregulated in the nuclear proteomic profile of HCT116-Dnmt1^Δ3-5^ cells indicate changes in Vimentin, E-Cadherin, which have been previously observed in this cell line [22, 28, 29]. We also show an increase in abundance of the EMT transcriptional driver Snail1, known to repress transcription of E-Cadherin, in contrast to findings published by [22]. In addition to this HCT116-Dnmt1^Δ3-5^ cells show decreased overall protein abundance of Beta-Catenin and redistribution of the protein to the cellular cytosol, in contradiction of previous research in this cell line by [22], who show that there is no change in Beta-Catenin protein expression, and that Beta-Catenin is redistributed to the nucleus. This redistribution of the protein is considered a contributing factor to the EMT phenotype and tumour reoccurrence in clinical models [31]. The EMT molecular signature observed in our study could not be recreated either by knockdown of endogenous WT protein and reintroduction of a WT DNMT1 construct into HCT116-Dnmt1^Δ3-5^ cells failed to repress Vimentin expression, indicating that the role of the mutant Dnmt1^Δ3-5^ protein in the induction of these EMT markers is independent of Dnmt1 protein levels.

Induction of EMT and dysregulation of proteasomal activity has been linked to the development of the CSC phenotype in a range of cancers, increased understanding of how this process is driven in malignancies will allow identifications of key EMT drivers and targeted development of novel therapeutic targets [32-35]. Progressive changes in methylation levels and patterns in CRC, as well as protein abundance changes in the key methyltransferase Dnmt1 have been well documented [7-10]. Understanding how the changes associated with Dnmt1-modulation of the epigenome and proteome during CRC progression may elucidate key mechanisms of EMT initiation within these cells and highlight how the EMT phenotype drives the development of CSC population. Finally, we show that in addition to investigations into Dnmt1s role as a methyltransferase, the hypomorphic cell line HCT116-Dnmt1^Δ3-5^ can be used to elucidate dysregulation of protein signalling networks in the nucleus, indicating that this model exhibits redistribution of Beta-Catenin as well as a molecular EMT phenotype across multiple signalling networks.

## MATERIALS AND METHODS

### Cell line culture and sample extraction

Colorectal cancer cell lines HCT116 and HCT116-Dnmt1^Δ3-5^ were regularly maintained in McCoy-5A media (Life Technologies, 16600-108, Carlsbad, CA) containing 10% fetal bovine serum (Life Technologies, 10438-026, Carlsbad, CA) and 1% streptomycin-penicillin (Life Technologies, 15140-148, Carlsbad, CA) at 37°C in CO2 incubator (5% CO2, 100% H2O). Cells were washed with warm PBS and then cells were detached using TrypLE express 1X (ThermoFisher). Detached cells were then washed with ice cold PBS, and lysed according to the following protocols.

### PCR analysis of Dnmt1 cDNA in HCT116 and HCT116-Dnmt1^Δ3-5^ cells

RNA was extracted from HCT116 and HCT116-Dnmt1^Δ3-5^ cells using TRIzol™ Reagent (Invitrogen) according to the manufactures protocol and stored at −80°C until required. Reverse transcriptase of RNA was perform using GoScript™ Reverse Transcription System (Promega) according to the manufacturers protocol. cDNA was stored at −20°C until analysed by polymerase chain reaction using GoTaq^®^G2 DNA polymerase (Promega) according to the manufacturers protocol. Annealing temperature was calculated on Tm of primer set (custom oligos, IDT) detailed in Table 1. PCR products were then analysed using gel electrophoresis, on a 0.9% agarose gel using GelRed^®^ Nucleic Acid Gel Stain (Biotium) and imaged using a G:BOX Ef imaging system (Syngene) and analysed using SnapGene software.

### Protein sample preparation

Cell pellets were re-suspended in RIPA buffer (50mM Tris-HCl pH8, 150mM NaCl, 0.1% SDS, 0.5% Deoxycholic acid, 1% NP-40.) supplemented with 1μl of 100x Protease inhibitor cocktail (ThermoFisher Scientific). Lysate was incubated for five minutes at room temperature before centrifugation at 16,000 x g at 4oC for 10 minutes (Eppendorf Centrifuge 5415R), supernatant was then removed and stored at −20°C or at −80°C for long term storage. Nuclear protein sample extract from washed cell pellets was carried out according to the manufactures protocol using cell fraction kit – standard (Abcam, ab109719). Nuclear pellets were resuspended in 100mM ammonium bicarbonate and lysed by sonication at 26 joules. Cell protein lysates were quantified using the manufactures protocol, PierceTM BCA protein assay kit (ThermoFisher Scientific). Colour change was read and protein quantifications calculated using PolarStar Omega (BMG Labtech) and Mars-data analysis software (BMG Labtech).

### SDS-PAGE & Immunoblotting

15μg of protein lysate, was incubated in a heating block (Benchmark) at 95oC for five minutes with Tris-glycine sample buffer (252mM Tris-HCl ph 6.8, 40% glycerol, 8% SDS, 0.01% bromophenol blue) and 1M DTT. Samples were then loaded and run on an 8% acrylamide gel (4ml of 2x gel buffer (160mM Tris-HCl pH7.4, 0.2M serine, 0.2M asparagine, 0.2M glycine),1.6ml of 40% acrylamide, 2.4ml H2O, 32µl 10% ammonium persulfate (APS), 16µl of tetramethylethylenediamine (TEMED)) with 1x Tris-glycine running buffer (25mM Tris, 192mM Glycine, 0.1% SDS). The gel was run for ten minutes at 100 volts (Biorad), then one hour and twenty minutes at 120 volts (Biorad) at 4oC. Proteins were transferred (XCell2 blot module, Invitrogen) to a nitrocellulose membrane (Amersham™ Protran™ 0.45μm NC) at 80 volts, 0.4 amps (Biorad) for three hours at 4oC in 1x Towbin transfer buffer (0.25M Tris pH8.6, 1.92M glycine, 0.025% SDS) using a XCell2 module (Invitrogen). Nitrocellulose membrane was then washed with 1xTBST twice and blocked for thirty minutes in 5% 1xTBST milk solution at room temperature. Membrane was then washed twice more in 1xTBST and incubated for two hours with primary antibody at 4oC on a rocker (Table 2). Membrane was then washed twice, for five minutes at room temperature, with 1xTBST, before incubating for one hour at room temperature with secondary LI-COR antibody (1xTBST 5% milk solution). Membrane was then washed three times for five minutes at room temperature with 1xTBST before imaging on a LI-COR Odyssey^®^ CLx and analysing with Image Studio Lite V5.2 software.

### Proteomic sample preparation and mass spectrometry

Nuclear pellets were lysed (0.1M TEAB, 0.1% SDS) with pulsed sonication, samples were centrifuged for 10 minutes, 13,000 x g, and supernatant protein quantified as described previously. Methanol/chloroform extraction was performed on 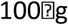 of protein for each lysate and pellet finally re-dissolved in 100ul of 6M Urea (Sigma), 2M thiourea (Sigma), 10mM HEPES buffer (Sigma), ph7.5. Samples were reduced (1M DTT), alkylated (5.5M iodoacetamide) then diluted in 400µl of 20mM ammonium biocarbonate (Sigma) and digested with trypsin (Promega) (1/50 w/w) overnight. Each fraction was acidified to <3.0 using trifluoroacetic acid (TFA) (Sigma) and solid phase extraction performed using Empore C18 96-well solid phase extraction plate (Sigma). Samples were eluted in 150µl of 80% acetonitrile, 0.5% acetic acid, lyophilised and then stored at −20C until re-suspension in 10µl 98% dH2O/acetonitrile and 0.1% formic acid with an internal quantification standard of 100 fmol of Waters enolase (Saccharomyces cerevisiae) and Hi3 Waters Escherichia coli standard added to each sample prior to use. Nano-LC separation was performed using the nanoAcquity UPLC 2G Trap column system (Waters) at a rate of 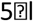 per min and washed with buffer A (98% dH2O/acetonitrile + 0.1% formic acid) for 5 minutes. Peptides were separated on an 50cm Acquity UPLC Peptide BEH C18 column (130Å, 1.7µm, 2.1mm x 150mm, 1/pkg (Waters)) over a 140 minute linear gradient of 5% to 40% with a flow rate of 0.3µl/min with buffer B (80% acetonitrile/dH2O + 0.1% formic acid) and completed with a 5 minutes rinse with 85% buffer B, at a flow rate of 300nl/min. Mass Spectrometry analysis was performed on a Waters Synapt G2Si HDMS system. Samples were sprayed directly using positive mode-ESI, and data was collected in MSE acquisition mode, alternating between low (5v) and high (20-40V) energy scans. Glu-fibrinopeptide (m/z = 785.8426, 100 fmol/µl) was used as LockMass, and was infused at 300 nl/min and sampled every 13 seconds for calibration. Raw data files were processed using ProteinLynx Global Server (PLGS, Waters) version 3.0.2, and optimal processing parameters were selected by Threshold Inspector software (Waters). Data-base search (Human Uniprot reference database July 2017 + yeast enolase P00924) was carried out using the Ion-accounting algorithm in PLGS (Waters), parameters included FDR of 1%, a maximum protein size of 500,000 Da, and trypsin cleavage with an allowable error of 1 missed cleavage with oxidation of methionine, and carbamidomethylation of cysteine modifications. Each sample replicate was merged into a single file to include the protein accession number (UniProt), protein description, and femtomoles on column. For protein quantification, the Hi3 calculation method of using the top 3 most intense tryptic peptides of both the internal standards and of each protein was used.

### Gene Ontology and protein interaction network analyses

Protein abundance (femtomoles on column) was used to identify significantly differential proteins (Student’s T-test) log2 ratio (hypomorph: wild-type) was computed. Graphical analysis of normalized mass spectrometry nuclear proteomic profiles was analysed using the R package ggplot2 and Morpheus GUI (https://software.broadinstitute.org/morpheus). Gene Ontology (GO) term enrichment was computed for the sets of significant (p<0.05) proteins using the Pathway Studio database (Elsevier), version 9.0. The most significantly differential GO terms were identified using the Rank Mobility Index for ranked terms between the hypomorph and wild-type datasets. Pathway Studio database network nodes and edges were supplemented by addition of STRING database interactions. Differentially abundant proteins were analysed using String DB, interaction networks were modified using internal settings for evidence-based interaction sources, and interactions represented based on their molecular action [36]. Interaction networks were than manually adapted to represent differential protein abundance as shown in nuclear proteomic profiling data of HCT116 and HCT116-Dnmt1^Δ3-5^ cells.

### Immunofluorescence microscopy

Cells were grown in 6 well plates (corning) and cell fixed using 4% paraformaldehyde and quenched using 50mM NH4Cl2. Cells were then permeabilise with 0.1% Triton100 and blocked in 10% normal goat serum for 30 minutes. Slides were then incubated with primary antibodies overnight at a dilution of 1:100 (see table 2 for antibody details). Slides were then incubated with secondary antibodies for 30 minutes at a dilution of 1:15,000 (Anti-Rabbit IgG 488 (Sigma SAB4600234) and Anti-Mouse IgG 555 (SAB4600068)). Coverslips were then mounted overnight at 4oC using ProLongTM Gold antifade mountant with DAPI (ThermoFisher P36931). Slides were then visualized using a Leica TCS SP8 microscope, aquired using LAS X Core software (V3.3.0) and analsyed using Fiji (released May 2017) with MBF “ImageJ for Microscopy” plugin (Dec 2009 release).

### Generation of shRNA-containing lentivirus

One million Hek293 FT cells were plated per well in a 6 well culture plate (Corning) and transfected 2 hours post-seeding with a 4-2-1 ratio of Plasmid: gagpol:vsvg, PEI transfection reagent (Sigma) in 100µl serum-free DMEM (LifeTech). Cells were transfected overnight, and media replaced the following morning with 2ml DMEM, 10%FCS, and 5% PenStrep. Cells were incubate for a further 24 hours before lentiviral-containing media was removed and filtered through a 0.45µm filter, lentiviral-containing media was then stored at −80°C until use.

### Transduction of cells with shRNA lentivirus

1 x 10^5^ HCT116 cells were plated per well in a 6 well culture plate (Corning), one cells have adhered remove culture medium and replace with diluted lentiviral media (500µl lentiviral media, 1.5ml DMEM, 10% FCS, 5% PenStrep, 5-10µg Polybrene (Sigma)). Cells were incubated in diluted lentiviral-containing media overnight, then grown for a further 24 hours in normal culture medium (DMEM, 10% FCS, 5% PenStep). Transient knockdown cell lines were lysed and protein samples extracted, stable cell lines were selected by 2µg/ml puromycin selection over five days before extraction of protein samples.

### Dnmt1 recovery experiment

Transfection of HCT116 and HCT116-Dnmt1^Δ3-5^ cells was performed using Lipofectamin 3000 Reagent (ThermoFisher) according to manufacturer’s protocol, transfection culture media was removed from cells 4 hours following transfection, and replaced with fresh complete culture medium. HCT116-Dnmt1^Δ3-5^ cells were transfected with pcDNA3 vector containing WT full length DNMT1 (36939, AddGene) and empty pcDNA3 vector as a transfection control (10792, AddGene). 48 hours following transfection cells were lysed and analysed by western blot.

## DECLARATIONS

### Acknowledgements

We thank Dr Mark Coldwell and members of his laboratory for assistance with cell-based assays.

### Authors’ contributions

Conceived and Designed Study: EB, RE, PS

Performed Experiments: EB, AS, PS, ND

Analysed Data: EB, AS, AL, JB, CB, RE

Wrote Paper: EB, RE, PS

Contributed Reagents/Materials: RE, PS, ND, ZW, YW

All authors read and approved the final manuscript

### Availability of data and material

The mass-spectrometry datasets generated and analysed during the current study are available in the PRIDE repository

### Ethics approval and consent to participate

N/A

## References

1. Clements EG, Mohammad HP, Leadem BR, Easwaran H, Cai Y, Van Neste L, et al. DNMT1 modulates gene expression without its catalytic activity partially through its interactions with histone-modifying enzymes. Nucleic Acids Res. 2012;40:4334–46.

2. Lei H, Oh SP, Okano M, Jüttermann R, Goss KA, Jaenisch R, et al. De novo DNA cytosine methyltransferase activities in mouse embryonic stem cells. Dev Camb Engl. 1996;122:3195–205.

3. Robertson KD. DNA methylation, methyltransferases, and cancer. Oncogene. 2001;20:3139–55.

4. Kanai Y, Ushijima S, Nakanishi Y, Sakamoto M, Hirohashi S. Mutation of the DNA methyltransferase (DNMT) 1 gene in human colorectal cancers. Cancer Lett. 2003;192:75–82.

5. Chen T, Hevi S, Gay F, Tsujimoto N, He T, Zhang B, et al. Complete inactivation of DNMT1 leads to mitotic catastrophe in human cancer cells. Nat Genet. 2007;39:391–6.

6. Sen GL, Reuter JA, Webster DE, Zhu L, Khavari PA. DNMT1 maintains progenitor function in self-renewing somatic tissue. Nature. 2010;463:563–7.

7. el-Deiry WS, Nelkin BD, Celano P, Yen RW, Falco JP, Hamilton SR, et al. High expression of the DNA methyltransferase gene characterizes human neoplastic cells and progression stages of colon cancer. Proc Natl Acad Sci U S A. 1991;88:3470–4.

8. Pathania R, Ramachandran S, Elangovan S, Padia R, Yang P, Cinghu S, et al. DNMT1 is essential for mammary and cancer stem cell maintenance and tumorigenesis. Nat Commun. 2015;6:6910.

9. Qu Y, Mu G, Wu Y, Dai X, Zhou F, Xu X, et al. Overexpression of DNA Methyltransferases 1, 3a, and 3b Significantly Correlates With Retinoblastoma Tumorigenesis. Am J Clin Pathol. 2010;134:826–34.

10. Zagorac S, Alcala S, Fernandez Bayon G, Bou Kheir T, Schoenhals M, González-Neira A, et al. DNMT1 Inhibition Reprograms Pancreatic Cancer Stem Cells via Upregulation of the miR-17-92 Cluster. Cancer Res. 2016;76:4546–58.

11. Rhee I, Bachman KE, Park BH, Jair K-W, Yen R-WC, Schuebel KE, et al. DNMT1 and DNMT3b cooperate to silence genes in human cancer cells. Nature. 2002;416:552–6.

12. Vertino PM, Yen RW, Gao J, Baylin SB. De novo methylation of CpG island sequences in human fibroblasts overexpressing DNA (cytosine-5-)-methyltransferase. Mol Cell Biol. 1996;16:4555–65.

13. Achour M, Fuhrmann G, Alhosin M, Rondé P, Chataigneau T, Mousli M, et al. UHRF1 recruits the histone acetyltransferase Tip60 and controls its expression and activity. Biochem Biophys Res Commun. 2009;390:523–8.

14. Cai Y, Geutjes E-J, de Lint K, Roepman P, Bruurs L, Yu L-R, et al. The NuRD complex cooperates with DNMTs to maintain silencing of key colorectal tumor suppressor genes. Oncogene. 2014;33:2157–68.

15. Song J, Du Z, Ravasz M, Dong B, Wang Z, Ewing RM. A protein interaction between beta-catenin and Dnmt1 regulates Wnt Signaling and DNA methylation in colorectal cancer cells. Mol Cancer Res. 2015;:molcanres.0644.2013.

16. Schermelleh L, Spada F, Easwaran HP, Zolghadr K, Margot JB, Cardoso MC, et al. Trapped in action: direct visualization of DNA methyltransferase activity in living cells. Nat Methods. 2005;2:751–6.

17. Stresemann C, Lyko F. Modes of action of the DNA methyltransferase inhibitors azacytidine and decitabine. Int J Cancer. 2008;123:8–13.

18. Rhee I, Jair KW, Yen RW, Lengauer C, Herman JG, Kinzler KW, et al. CpG methylation is maintained in human cancer cells lacking DNMT1. Nature. 2000;404:1003–7.

19. Egger G, Jeong S, Escobar SG, Cortez CC, Li TWH, Saito Y, et al. Identification of DNMT1 (DNA methyltransferase 1) hypomorphs in somatic knockouts suggests an essential role for DNMT1 in cell survival. Proc Natl Acad Sci U S A. 2006;103:14080–5.

20. Spada F, Haemmer A, Kuch D, Rothbauer U, Schermelleh L, Kremmer E, et al. DNMT1 but not its interaction with the replication machinery is required for maintenance of DNA methylation in human cells. J Cell Biol. 2007;176:565–71.

21. Cai Y, Tsai H-C, Yen R-WC, Zhang YW, Kong X, Wang W, et al. Critical threshold levels of DNA methyltransferase 1 are required to maintain DNA methylation across the genome in human cancer cells. Genome Res. 2017;27:533–44.

22. Espada J, Peinado H, Lopez-Serra L, Setién F, Lopez-Serra P, Portela A, et al. Regulation of SNAIL1 and E-cadherin function by DNMT1 in a DNA methylation-independent context. Nucleic Acids Res. 2011;39:9194–205.

23. Du Z, Song J, Wang Y, Zhao Y, Guda K, Yang S, et al. DNMT1 stability is regulated by proteins coordinating deubiquitination and acetylation-driven ubiquitination. Sci Signal. 2010;3:ra80.

24. Komaba S, Coluccio LM. Localization of myosin 1b to actin protrusions requires phosphoinositide binding. J Biol Chem. 2010;285:27686–93.

25. Lamouille S, Xu J, Derynck R. Molecular mechanisms of epithelial-mesenchymal transition. Nat Rev Mol Cell Biol. 2014;15:178–96.

26. Hanahan D, Weinberg RA. Hallmarks of cancer: the next generation. Cell. 2011;144:646–74.

27. Pino MS, Kikuchi H, Zeng M, Herraiz M-T, Sperduti I, Berger D, et al. The epithelial to mesenchymal transition is impaired in colon cancer cells with microsatellite instability. Gastroenterology. 2010;138:1406–17.

28. Wang H, Wang H-S, Zhou B-H, Li C-L, Zhang F, Wang X-F, et al. Epithelial–Mesenchymal Transition (EMT) Induced by TNF-α Requires AKT/GSK-3β-Mediated Stabilization of Snail in Colorectal Cancer. PLOS ONE. 2013;8:e56664.

29. Wheelock MJ, Shintani Y, Maeda M, Fukumoto Y, Johnson KR. Cadherin switching. J Cell Sci. 2008;121 Pt 6:727–35.

30. Peng L, Yuan Z, Ling H, Fukasawa K, Robertson K, Olashaw N, et al. SIRT1 deacetylates the DNA methyltransferase 1 (DNMT1) protein and alters its activities. Mol Cell Biol. 2011;31:4720–34.

31. Holthoff ER, Spencer H, Kelly T, Post SR, Quick CM. Pathologic features of aggressive vulvar carcinoma are associated with an epithelial-mesenchymal transition. Hum Pathol. 2016;56:22–30.

32. Banno A, Garcia DA, van Baarsel ED, Metz PJ, Fisch K, Widjaja CE, et al. Downregulation of 26S proteasome catalytic activity promotes epithelial-mesenchymal transition. Oncotarget. 2016;7:21527–41.

33. Fan F, Samuel S, Evans KW, Lu J, Xia L, Zhou Y, et al. Overexpression of snail induces epithelial-mesenchymal transition and a cancer stem cell-like phenotype in human colorectal cancer cells. Cancer Med. 2012;1:5–16.

34. Mani SA, Guo W, Liao M-J, Eaton EN, Ayyanan A, Zhou AY, et al. The epithelial-mesenchymal transition generates cells with properties of stem cells. Cell. 2008;133:704–15.

35. Zhu P, Zhao N, Sheng D, Hou J, Hao C, Yang X, et al. Inhibition of Growth and Metastasis of Colon Cancer by Delivering 5-Fluorouracil-loaded Pluronic P85 Copolymer Micelles. Sci Rep. 2016;6:20896.

36. Szklarczyk D, Gable AL, Lyon D, Junge A, Wyder S, Huerta-Cepas J, et al. STRING v11: protein-protein association networks with increased coverage, supporting functional discovery in genome-wide experimental datasets. Nucleic Acids Res. 2019;47:D607–13.

